# Expanding the watch list for potential Ebola virus antibody escape mutations

**DOI:** 10.1101/516161

**Authors:** Jagdish Suresh Patel, Caleb J. Quates, Erin L. Johnson, F. Marty Ytreberg

## Abstract

The 2014 outbreak of Ebola virus (EBOV) in Western Africa is the largest recorded filovirus disease outbreak and lead to the death of over 11,000 people. This deadly virus still poses a grave epidemic threat as evidenced by the current (since May 2018) EBOV outbreak in the Democratic Republic of the Congo which has already claimed the lives of over 250 people. One important strategy for combating EBOV epidemics is to anticipate how the evolution of EBOV might undermine treatment since the development of vaccines and antibody therapies are typically based on a single strain (often the 1976 Mayinga) of the EBOV envelope glycoprotein (GP). In a previous study we initiated a watch list of potential antibody escape mutations of EBOV by modeling interactions between EBOV GP and the monoclonal antibody KZ52. This watch list was generated using molecular modeling to estimate the effect of every possible mutation of GP. The final watch list containing 34 mutations were predicted to disrupt GP-KZ52 binding but not to disrupt the ability of GP to fold and to form trimers. In this study, we expand our watch list by including three more monoclonal antibodies with distinct epitopes on GP, namely Antibody 100 (Ab100), Antibody 114 (Ab114) and 13F6-1-2. Our updated watch list contains 127 mutations, three of which have been seen in humans or are experimentally associated with reduced efficacy of antibody treatment. We believe mutations on this broad watch list require attention since they may be a signal of an evolutionary response from EBOV to treatment that could diminish the effectiveness of interventions.

## Introduction

Ebola virus (EBOV) has five known species: Zaire EBOV, Sudan EBOV, Ta Forest EBOV, Bundibugyo EBOV, and Reston EBOV and can cause disease with up to 90% case fatality rates. [1, 2] The outbreak of Zaire EBOV infection in Western Africa between 2014 to 2016 is the largest recorded filovirus disease outbreak with death typically occurring 5-7 days after the appearance of symptoms.[3] This outbreak receded in 2016, but EBOV still constitutes an epidemic threat as evidenced by the current (since May 2018) Zaire EBOV outbreak in the Democratic Republic of the Congo which has already claimed lives of 271 people and more than 458 positive cases (WHO report, 4^th^ December 2018).[4]

The 2014–2016 EBOV outbreak in West Africa led to clinical development of therapeutics and vaccines.[5, 6] The success of ZMapp, a cocktail of three chimeric monoclonal antibodies (mAbs) derived from immunized mice, in nonhuman primates (NHP) demonstrated the potential of mAb therapies against EBOV infection, and ZMapp is currently undergoing human trials.[7–9] Subsequently, several other mAbs were designed and demonstrated protection against EBOV infection and some of them are currently being used to manage the current EBOV outbreak. (https://www.nih.gov/news-events/news-releases/nih-begins-testing-ebola-treatment-early-stage-trial)[10, 11] To prepare for future outbreaks it is critical to anticipate and monitor EBOV evolution since it could lead to antibody escape mutants that could compromise treatment efforts. Sequencing studies have revealed that there is a significant genetic variation in EBOV. Most of the variation is found in the EBOV glycoprotein (GP) that is the target for the all mAbs currently under development and being used in the management of the current EBOV outbreak.[12–14]

To address the potential threat of EBOV evolution outpacing antibody treatment efforts we previously initiated a watch list of potential antibody escape mutants for the EBOV GP.[15] We focused on the KZ52 mAb as it had a 3-D experimental structure bound to EBOV GP available. In this previous study, we combined use of FoldX software with molecular dynamics simulations to estimate folding and binding stabilities. We placed 34 mutations on the watch list by considering every possible EBOV GP mutation and choosing those that disrupt binding between GP and KZ52 but do not disrupt the ability of GP to fold and bind to form a complex. One of these 34 mutations (*N550K*) was seen in humans in a previous outbreak.

The aim of this study is to expand our watch list of potential antibody escape mutants for EBOV GP. After we published our previous watch list, three more 3-D structures of mAbs interacting with different epitopes on EBOV GP (necessary for our molecular modeling approach) were published in the Protein Data Bank (PDB). In this study, we expand our watch list to include possible antibody escape mutations from three antibodies with distinct epitopes: Antibody 100 (Ab100), Antibody 114 (Ab114) and 13F6-1-2. Our watch list now contains 127 mutations, three of which have been seen in humans or are experimentally associated with reduced efficacy of antibody treatment.

## Methods

For a mutation to be placed on a watch list for EBOV it must: (1) disrupt binding to a protective antibody, and (2) leave the viral proteins functional. It is thus necessary to determine how amino acid mutations alter stabilities (ΔΔ*G* values) for GP folding, forming a trimer and binding to the antibody. In our previous work, we have obtained ΔΔ*G* values of GP folding, GP trimer formation and GP binding to KZ52 antibody using our molecular dynamics (MD) plus FoldX approach.[15] In this study we calculate ΔΔ*G* values of binding for Ab100, Ab114 and 13F6-1-2 antigen-antibody complexes using the same modeling approach.

### Structure Preparation EBOV GP – Ab100 and Ab114 complexes

Structures of Ab100 and Ab114 bound to EBOV GP were obtained from the Protein Data Bank (PDB) accession number 5FHC.[16] The EBOV GP amino acid sequence was based on the 1976 Mayinga strain, just like our previous study. The PDB file 5hfc.pdb contains coordinates of both Ab100 and Ab114 bound to EBOV GP (GP1 & GP2) monomer at different sites. This file was first modified to remove all but one copy each of GP1, GP2, antibody light chain and antibody heavy chain of Ab100 and Ab114 (one third of the GP-Ab100/Ab114 trimeric complex), and then was split into two files where one file had the GP – Ab100 complex and the other had the GP – Ab114 complex. The MODELLER software[17] was then used to build the missing residues. This included residues 190 – 213 that are predicted to be intrinsically disordered; the resulting complexes had no secondary structure content in this region. The full EBOV trimer protein complex was then created using the symexp command in PyMOL (see Figs 1A and 2A) and contained three copies each of GP1 (residues 33 - 278), GP2 (residues 502 – 599), heavy and light chains of the antibody.

**Fig 1.**
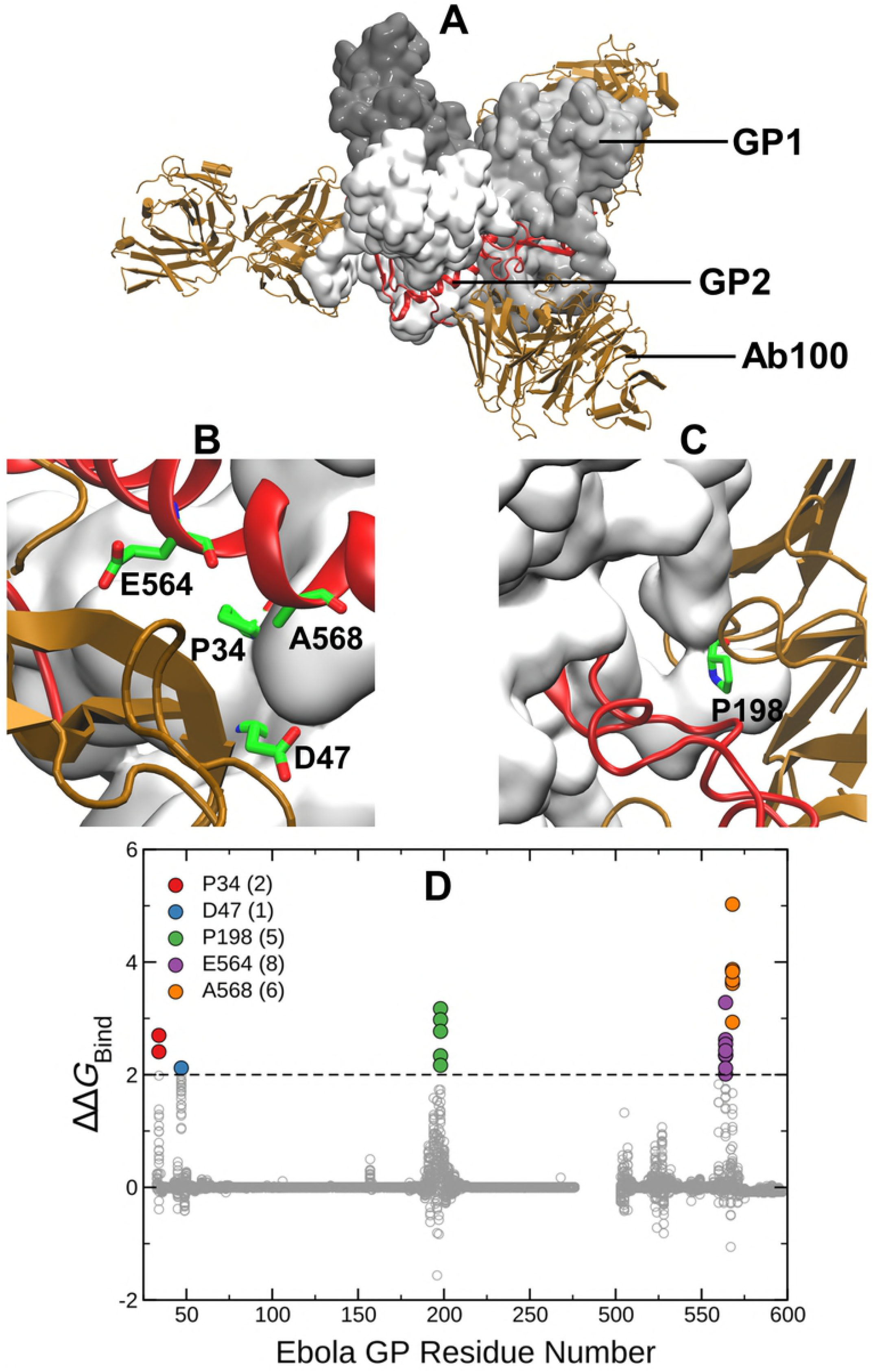
(A) Structure of EBOV GP trimer in complex with the Ab100 antibody. GP1 is gray, GP2 is red and the heavy and the light chains of Ab100 are brown. (B) & (C) Mutations in the GP complex with ΔΔ*G*_Bind_ > 2 kcal/mol (i.e., above black dashed line) are considered disruptive and are highlighted using green stick representation. (D) ΔΔ*G*_Bind_ values (gray circles) for all 19 possible mutations at each site of GP1 (33 – 278) and GP2 (502 – 599). Different colors and counts in the legend indicate locations and number of mutations with ΔΔ*G*_Bind_ > 2 kcal/mol on the GP complex.

**Fig 2.**
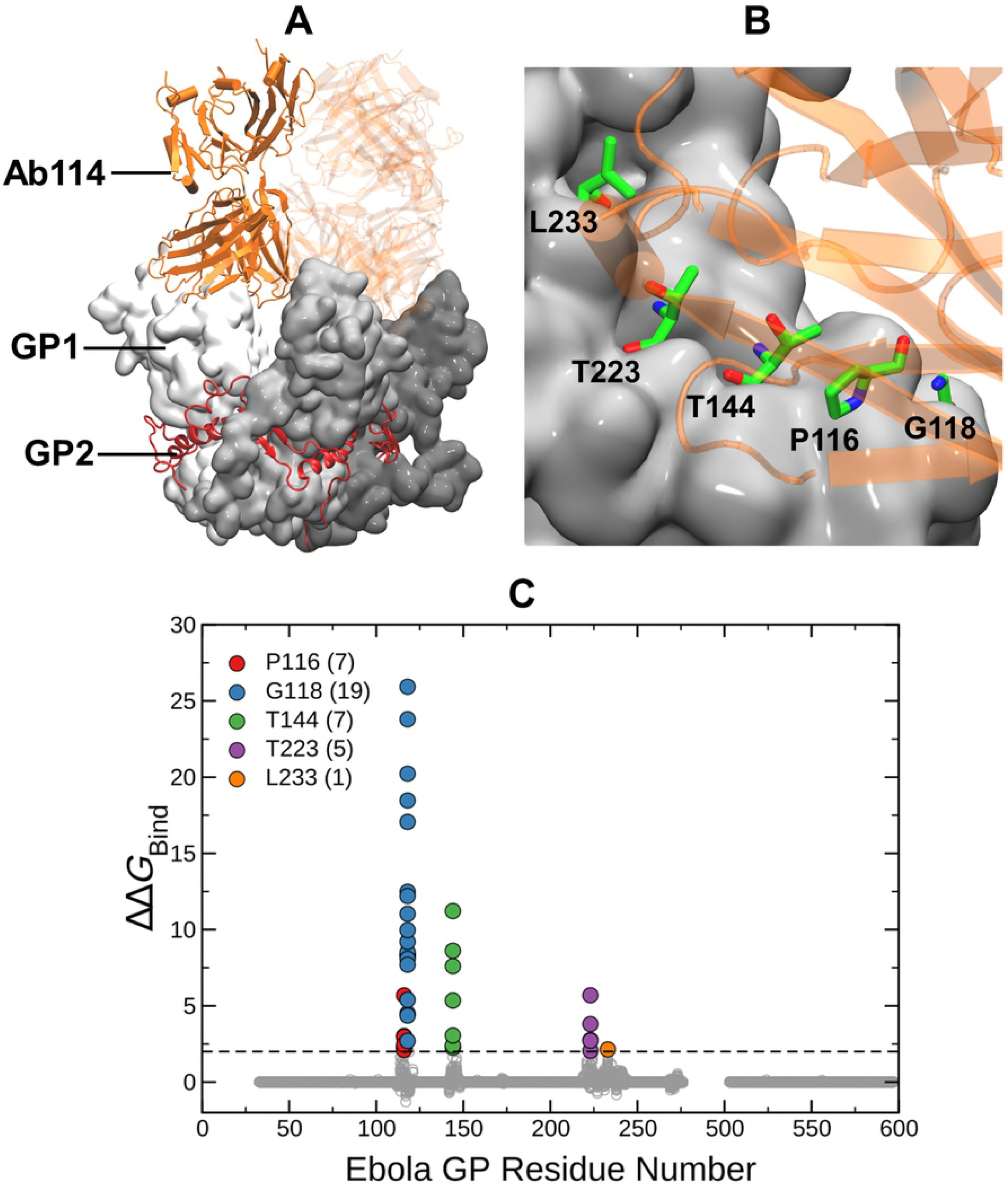
(A) Structure of EBOV GP trimer in complex with the Ab114 antibody. GP1 is gray, GP2 is yellow and the heavy and the light chains of Ab114 are orange. (B) Mutations in the GP complex with ΔΔ*G*_Bind_ > 2 kcal/mol (i.e., above black dashed line) are considered disruptive and are highlighted using green stick representation. (D) ΔΔ*G*_Bind_ values (gray circles) for all 19 possible mutations at each site of GP1 (33 – 278) and GP2 (502 – 599). Different colors and counts in the legend indicate locations and number of mutations with ΔΔ*G*_Bind_ > 2 kcal/mol on the GP complex.

### EBOV mucin-like domain peptide – 13F6-1-2 antibody complex

Crystal structure of the 13F6-1-2 Fab fragment bound to its EBOV GP mucin-like domain (GP MLD) peptide epitope (11 amino acids, VEQHHRRTDND) was downloaded from the PDB using the 2QHR accession number.[18] Unlike other epitopes used in our previous and current study, this structure was based on the 1976 Eckron strain. However, the alignment of the 11-residue long peptide was 100% identical to the 1976 Mayinga strain. The PDB file 2qhr.pdb was edited to remove everything except GP MLD peptide (residues 404 – 414) and heavy and light chains of 13F6-1-2 antibody. (see Fig 3A) WHAT IF web server (https://swift.cmbi.umcn.nl) was then used to add the missing atoms to the complex structure.

**Fig 3.**
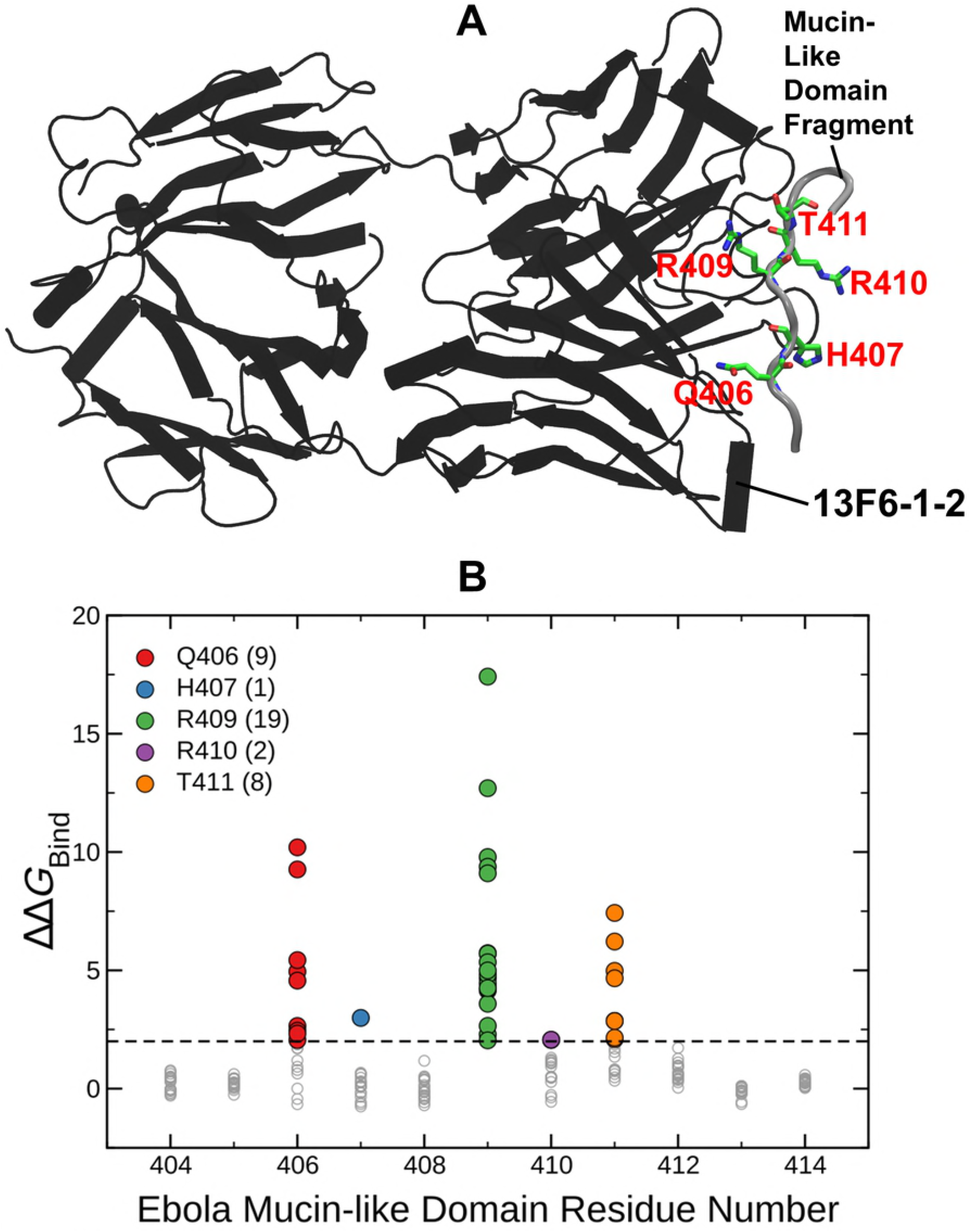
(A) Structure of EBOV GP MLD peptide bound to the 13F6-1-2 antibody. GP MLD peptide is in gray tube representation, and the heavy and the light chains of 13F6-1-2 are black. Mutations in the GP MLD peptide with ΔΔ*G*_Bind_ > 2 kcal/mol (i.e., above black dashed line) are considered disruptive and are highlighted using green stick representation. (B) ΔΔ*G*_Bind_ values (gray circles) for all 19 possible mutations at each of the 11 sites of GP MLD peptide (404 –414). Different colors and counts in the legend indicate locations and number of mutations with ΔΔ*G*_Bind_ >2 kcal/mol on the GP complex.

### Molecular Dynamics Simulations

All three EBOV – antibody complexes were subjected to atomistic MD simulations using the protocol reported in our previous study. Briefly, the software package GROMACS 5.0.7[19] was used for all MD simulations with the Charmm22* forcefield.[20] GP MLD peptide – 13F6-1-2 simulations were 100 ns, and the GP – Ab100 and Ab114 production simulations were run for a shorter 50 ns due to the large number of atoms in the final simulation box. During the production simulation snapshots were saved every 1 ns resulting in 50 snapshots for each of the GP – Ab100 and GP – Ab114 systems, and 100 snapshots for the peptide – 13F6-1-2 complex.

### FoldX

Snapshots of all three complexes were analyzed using FoldX software.[21, 22] As with our previous study, we initially minimize each snapshot six times in succession using the RepairPDB command to obtain convergence of the potential energy. BuildModel command was then used to generate all possible 19 single mutations in EBOV GP and GP MLD peptide at each amino acid site. Finally, the binding stability of the protein complex due to each mutation was estimated using AnalyseComplex command. For each mutation, we then estimated ΔΔ*G*_Bind_ by averaging the FoldX results across all individual snapshot estimates.

To calculate ΔΔ*G*_Bind_ values for all possible 19 mutations at amino acid site of EBOV GP and GP MLD peptide, we carried out 980,400 FoldX calculations (344 GP residues × 19 possible mutations at each site × 50 MD snapshots × 3 copies of GP-Antibody in a complex) for each GP – Ab100 and GP – Ab114 complexes and 20,900 calculations (11-residue MLD peptide × 19 possible mutations × 100 MD snapshots) for GP MLD peptide – 13F6-1-2 antibody complex. Averaging estimates across all individual snapshots ultimately resulted in 6,441 ΔΔ*G*_Bind_ values each for Ab100 and Ab114 antibody complexes and 209 ΔΔ*G*_Bind_ values for 13F6-1-2 antibody complex. (see SI Files)

## Results and discussion

We have expanded the watch list generated by us in a previous study by including antibody escape mutations against three additional antibodies interacting with EBOV GP. These watch list mutations are those predicted to both disrupt GP - antibody binding and yet allow GP to fold and form trimers.

The EBOV GP is a class I fusion protein consisting of disulfide-linked subunits, GP1 and GP2, that bind to form a chalice-shaped trimer. (see Figs 1A & 2A) Ab100 interacts at the base of the GP trimer (see Fig 1A). This interaction is similar to that of KZ52, the prototypic neutralizing antibody used in our previous study. However, Ab100 contacts GP1 and GP2 of a monomer and the disordered (residue 190 - 213) loop of the neighboring monomer (see Fig 1A) in contrast to KZ52 which interacts with GP2 of a single monomer. The epitope for Ab114 spans both the glycan cap and the inner chalice of GP, (see Fig 2A) where it remains bound after proteolytic cleavage of the glycan cap and prevents interaction of cleaved GP to its host receptor.[16] 13F6-1-2 is a monoclonal antibody that binds to an 11-residue peptide located in the heavily glycosylated mucin-like domain (MLD) of the EBOV GP.[18] (see Fig 3A)

Fig 4 is a graphical representation of our expanded watch list of EBOV mutations which includes KZ52, Ab100, Ab114 and 13F6-1-2 antibody complexes. The maximum functional stability change (i.e., folding and trimer formation) for all mutations is plotted against the corresponding change in the antibody binding affinity. The 127 mutations in the lower right quadrant are those that belong on the watch list since they are classified as both functional and disruptive to antibody binding. There were 21, 33, 39 and 34 mutations respectively in Ab100, Ab114, 13F6-1-2 and KZ52 antibody complexes. The specific mutations on the watch list are given in the Table 1 and shows that they are concentrated at just six residues in KZ52 and at five residues in each of the other three antibody complexes. (see Figs 1B & 1C, Fig 2B and Fig 3A) All of these mutations lie at the interface as one might intuitively expect from the structure, however, most mutations at the interface are not predicted to disrupt antibody binding.

**Fig 4.**
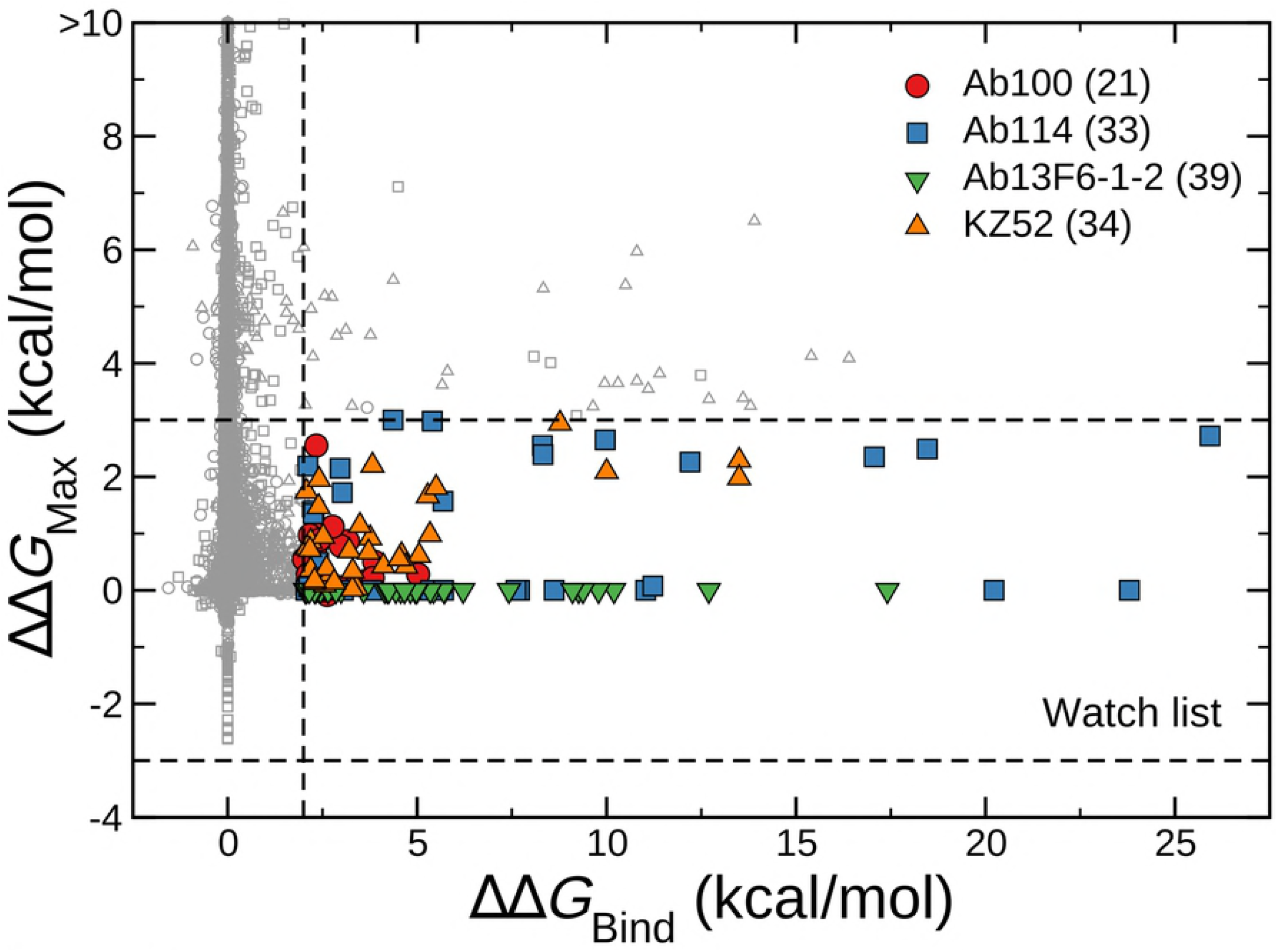
Maximum of folding stability, dimer binding stability (interaction of GP1 and GP2) or trimer binding stability (interaction of a GP1-GP2 dimer with other dimers), ΔΔ*G*_Max_, as a function of ΔΔ*G*_Bind_ for all antibody complexes. ΔΔGMax values are considered to be zero for the intrinsically disordered 11-residue MLD peptide. Symbols in the inset legend indicate the corresponding antibody. Watch list mutations are shown as colored symbols and are predicted to disrupt binding to any one of the four antibodies (KZ52, Ab100, Ab114 and 13F6-1-2) but not to disrupt GP folding and trimer formation. Consistent with our previous study, mutations with ΔΔ*G*_Bind_ > 2 kcal/mol are considered disruptive to antibody binding and those with –3 < ΔΔ*G*_Max_ < 3 kcal/mol are considered functional. The number of watch list mutations associated with each antibody is shown in the legend.

**Table 1.**
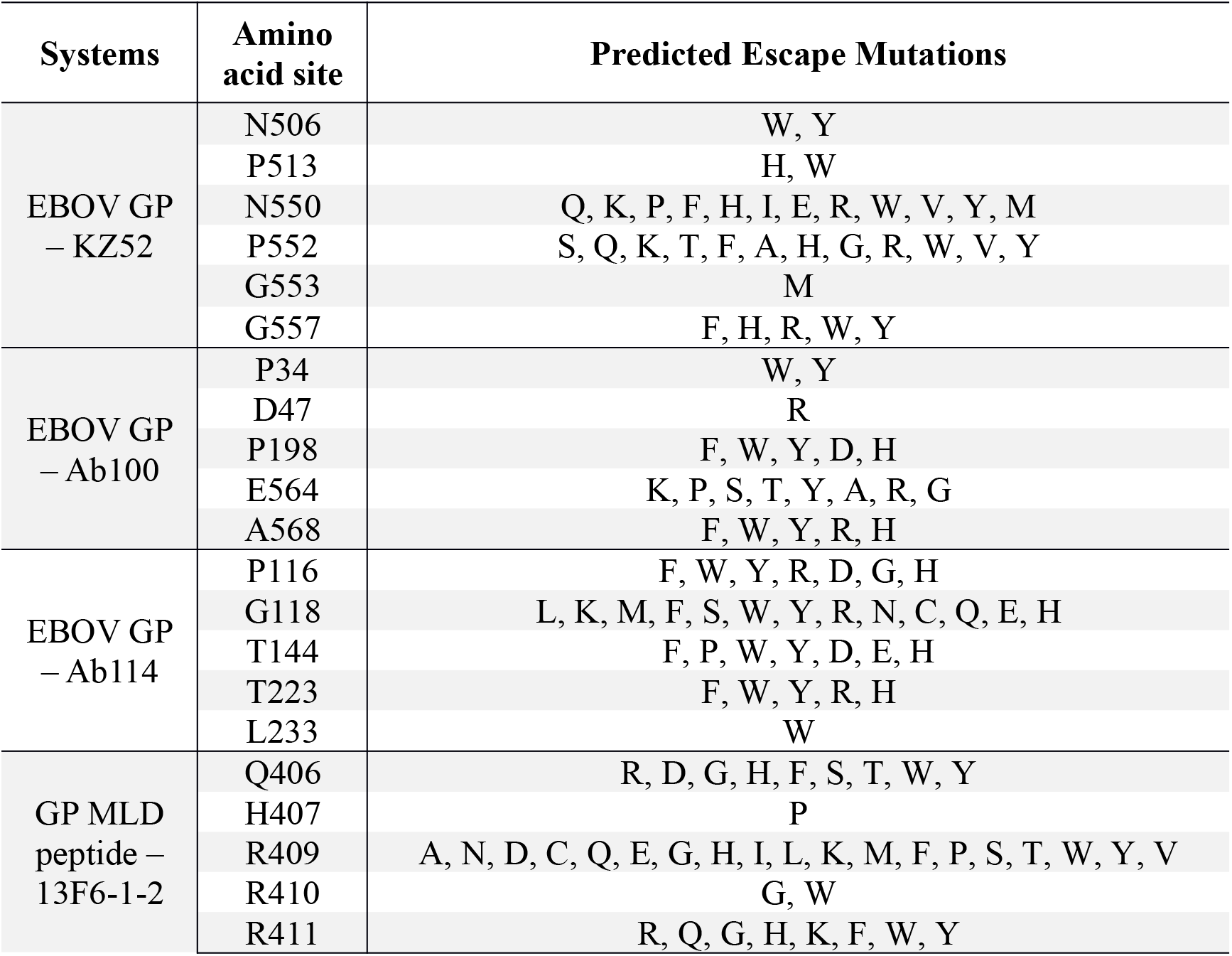
Watch list mutations. Watch list mutations are shown for all four antibodies (KZ52, Ab100, Ab114 and 13F6-1-2) at a predicted amino acid sites on EBOV GP.

| Systems | Amino acid site | Predicted Escape Mutations |
| --- | --- | --- |
| EBOV GP<br>– KZ52 | N506 | W, Y |
|  | P513 | H, W |
|  | N550 | Q, K, P, F, H, I, E, R, W, V, Y, M |
|  | P552 | S, Q, K, T, F, A, H, G, R, W, V, Y |
|  | G553 | M |
|  | G557 | F, H, R, W, Y |
| EBOV GP<br>– Ab100 | P34 | W, Y |
|  | D47 | R |
|  | P198 | F, W, Y, D, H |
|  | E564 | K, P, S, T, Y, A, R, G |
|  | A568 | F, W, Y, R, H |
| EBOV GP<br>– Ab114 | P116 | F, W, Y, R, D, G, H |
|  | G118 | L, K, M, F, S, W, Y, R, N, C, Q, E, H |
|  | T144 | F, P, W, Y, D, E, H |
|  | T223 | F, W, Y, R, H |
|  | L233 | W |
| GP MLD<br>peptide –<br>13F6-1-2 | Q406 | R, D, G, H, F, S, T, W, Y |
|  | H407 | P |
|  | R409 | A, N, D, C, Q, E, G, H, I, L, K, M, F, P, S, T, W, Y, V |
|  | R410 | G, W |
|  | R411 | R, Q, G, H, K, F, W, Y |

If any of the mutations in Table 1 were to appear in a real population, it would indicate a possible increased risk of escaping the normal immune response or render treatments with these antibodies ineffective. Mutation *N550K* against KZ52 has already appeared in humans and thought to have been infected by gorillas in Central Africa between 2001 and 2003. This mutation is present in all sequenced isolates from that outbreak.

Davidson et al.[23] conducted an alanine-scanning mutagenesis study on GP that can be qualitatively compared to our work. They individually mutated each residue of the GP protein to alanine and measured changes in GP-KZ52 binding affinities relative to the unmutated form. They identified five residues that are critical for KZ52 antibody binding: C511, N550, D552, G553, and C556. Three of these sites are found on our watch list in Table 1 (N550, D552, and G553) and 25 of the 34 (74%) watch list mutations are found at these three sites.

In the study of MB-003,[24] a plant-derived monoclonal antibody cocktail composed of c13C6, 13F6-1-2, and c6D8 used effectively in treatment of EBOV virus infection in non-human primates, was unable to protect two of six animals when initiated one- or two-days postinfection. Investigation of a mechanism of viral escape in one of the animals shown five nonsynonymous mutations in the monoclonal antibody target sites. Among these mutations *Q406R* and *R409C* were linked to a reduction in 13F6-1-2 antibody binding.[24] Both of these mutations were correctly identified by our modeling strategy and are present on the watch list.

In spite of our efforts to expand the watch list, it remains incomplete and putative for several reasons. The list covers only four epitopes for which experimental structures of antibodies interacting with viral proteins are available. However, there are other epitopes known for EBOV GP.[10] With more experimental structures it would be possible to further expand the watch list. The size of the watch list depends on how we define the threshold for antibody binding (Fig 4). If the threshold is lowered from 2.0 kcal/mol, the watch list will naturally grow in size. The reasoning behind using a different cutoff for functional as compared to antibody binding is described in our previous study. The watch list only includes mutations that are individually predicted to disrupt antibody binding while remaining functional, however, it is possible that immune escape could arise by the cumulative effect of several changes on antibody binding. How multiple substitutions interact to produce cumulative effects on stability is not well understood and is an important consideration for future studies. The watch list has not been experimentally validated either in terms of mutational effects on GP folding and binding affinities, nor on the downstream immune system consequences. Our hope is that this work will encourage such research.

In summary, we have expanded the watch list of potential antibody escape mutations of EBOV by including three more antibody complexes: EBOV GP – Ab100, GP – Ab114 and GP Mucin-like domain (GP MLD) peptide – 13F6-1-2. The watch list now contains 127 mutations in 21 sites in EBOV GP. Mutations from the watch list that appear during an outbreak deserve attention since they may be a signal of an evolutionary response from the virus that could reduce the efficacy of treatment efforts. Ab114 has been recently approved for the first time to treat infected individuals during the current EBOV outbreak in Democratic Republic of Congo. (https://www.nih.gov/news-events/news-releases/nih-begins-testing-ebola-treatment-early-stage-trial) We hope our watch list will serve as a useful reference for the public health and emerging infectious disease monitoring agencies. The watch list can still be expanded if more experimental structures of EBOV – antibody complexes become available. In fact, as we were preparing this manuscript, the crystal structure of mAb CA45 bound to GP1 and GP2 interface of EBOV GP were published by Janus et al.[25] Lastly, we believe our in-silico approach could be applied to determine watch lists for other viruses provided experimental structures are available and for future design and optimization efforts of antibodies.

## Acknowledgements

This research was supported by the Center for Modeling Complex Interactions sponsored by the NIGMS under award number P20 GM104420 and National Science Foundation EPSCoR Track-II under award number OIA1736253. Computer resources were provided in part by the Institute for Bioinformatics and Evolutionary Studies Computational Resources Core sponsored by the National Institutes of Health (P30 GM103324). This research also made use of the computational resources provided by the high-performance computing center at Idaho National Laboratory, which is supported by the Office of Nuclear Energy of the U.S. DOE and the Nuclear Science User Facilities under Contract No. DE-AC07-05ID14517. The funders had no role in study design, data collection and analysis, decision to publish, or preparation of the manuscript.

## Supporting information

**S1 File. S1_Ab100_data.xlsx.** Excel spreadsheet with estimated stability effects of all 6,441 mutations against Ab100.

**S2 File. S2_Ab114_data.xlsx.** Excel spreadsheet with estimated stability effects of all 6,441 mutations against Ab114.

**S3 File. S3_13F6-1-2_data.xlsx.** Excel spreadsheet with estimated stability effects of all 209 mutations against 13F6-1-2.

**S4 File. S4_expanded_watchlist.xlsx.** Excel spreadsheet with estimated binding stability effects of watch list mutations against all the four antibodies (KZ52, Ab100, Ab114, and 13F6-1-2).

